# Genetic lineage tracing identifies intermediate mesoderm as a novel contributor to mammalian kidney lymphatics

**DOI:** 10.1101/2025.07.18.665353

**Authors:** Daniyal J Jafree, Lauren G Russell, Athanasia Stathopoulou, Christopher J Rowan, Andrew T White, Charlotte O’Riordan, Maria Kolatsi-Joannou, Karen L Price, Sarah Ivins, Liam A Ridge, Catherine Roberts, Jennie C Chandler, Laura Wilson, Dale Moulding, Julie Siegenthaler, Adrian S Woolf, Paul R Riley, Christiana Ruhrberg, Peter J Scambler, Norman D Rosenblum, David A Long

**Author notes:** Corresponding author Professor David A Long, Professor in Paediatric Nephrology & Wellcome Trust Investigator in Science Developmental Biology and Research & Teaching Department, UCL Great Ormond Street Institute of Child Health 30 Guilford Street, London, WC1N 1EH, UK. D.J.J., L.G.R. and A.S. contributed equally to this work.

## Abstract

The lymphatic vasculature is essential for fluid homeostasis, immune regulation and possesses diverse organ-specific functions. During development, lymphatic endothelial cells (LEC) arise from multiple progenitor sources that form organ-specific lymphatic networks. While the origins of LECs in the heart, skin, and mesentery have been studied, those in the kidney remain unresolved. Here, we combined genetic lineage tracing in mouse embryos with optical clearing and high-resolution three-dimensional imaging to identify two distinct progenitor sources of kidney lymphatics. The majority of kidney LECs originate from a *Tie2⁺* endothelial lineage previously linked to venous or capillary vessels. Approximately 15% derive from *Osr1⁺* intermediate mesoderm, a lineage that generates kidney nephrons and stroma. *Osr1⁺-*derived LECs were absent from the heart, mesentery, and skin, indicating a kidney-specific contribution, and arose independently of nephron and stromal lineages. Both *Tie2⁺* and *Osr1⁺* lineages contributed to vessel sprouting and *de novo* formation of lymphatic clusters. Revealing a novel cellular origin of LECs and identifying a dual origin for kidney lymphatics, we demonstrate that *de novo* lymphatic formation can occur from both shared and organ-specific progenitors. This work advances our understanding of how lymphatics assemble during development and provides a framework for targeting kidney lymphatics in disease.

## INTRODUCTION

Lymphatic vessels are present in nearly all mammalian organs, where they maintain tissue fluid balance, transport macromolecules and regulate immune responses. During disease, lymphatics expand to facilitate the resolution of inflammation and tissue repair. Beyond these conserved roles, lymphatics exert organ-specific functions^1–3^, such as cerebrospinal fluid drainage from the central nervous system and nasopharynx^4–6^, maintenance of stem cells niches in the heart, skin and intestines^7–10^ and thermoregulation in adipose tissue^11,12^. However, it remains unclear how functional heterogeneity is established during development.

One emerging hypothesis is that the developmental origin of lymphatic endothelial cells (LEC), which line lymphatic vessels, underpins their organ-specific characteristics^13,14^. The differentiation of LECs from venous endothelium, followed by their proliferation and sprouting into lymphatic vessels, termed lymphangiogenesis, was long thought to be the primary mechanism of lymphatic development^15–20^. However, additional progenitor sources that give rise to LECs *de novo* have been identified^21,22^. A second model, termed lymphvasculogenesis, has been proposed, based on the presence of lymphatic clusters anatomically distinct from sprouting lymphatics within the developing meninges^23^, heart^24,25^, skin^26,27^, mesentery^28^ and kidney^29^. Demonstrated by live imaging studies in zebrafish embryos^30–32^, lymphangiogenesis and lymphvasculogenesis cooperate to generate organ-specific lymphatic networks.

Genetic lineage tracing studies in mice have begun to unravel the diversity of progenitors giving rise to LECs, with origins traced to the second heart field^25,33^, hemogenic endothelium^28,34^ and tissue-resident blood capillaries^27^. Moreover, it was shown that paraxial mesoderm gives rise to lymphatic vessels and isolated clusters in several different organs^35,36^. Despite these advances, lineage tracing studies have not yet established the developmental origins of kidney lymphatics. This gap is notable, given the kidney’s central role in physiological homeostasis, the expansion of lymphatics in kidney diseases, and the therapeutic interest in targeting lymphatics in conditions such as chronic kidney disease^37–40^, polycystic kidney disease^29,41,42^ and transplant rejection^43–45^.

We previously showed that kidney lymphatic development involves both lymphangiogenesis and lymphvasculogenesis^29^, but the identity of the contributing progenitors remained undefined. Within the mouse metanephros, the direct precursor to the adult kidney, intermediate mesoderm gives rise to epithelial, stromal and blood vasculature^46^, and is therefore a plausible source of kidney lymphatics. However, whether LECs within the kidney arise from this lineage, or instead share progenitors with lymphatics in other organs, has not been established.

To address this, we combined genetic lineage tracing, optical clearing, and three-dimensional (3D) imaging^45,47^ in mouse embryos to quantitatively map the origins of kidney LECs. We show that the majority of kidney lymphatics arise from *Tie2⁺* progenitors, an origin shared by lymphatics in other organs^20,36^. Conversely, 15% of LECs arise from *Osr1⁺* intermediate mesoderm, distinct from the lineage composition of lymphatics in other organs. These findings reveal that kidney lymphatics originate from complementary, developmentally distinct sources, refining our understanding of organ-specific lymphatic development and the broader lineage architecture of the kidney.

## RESULTS

### The majority of kidney lymphatics originate from endothelial progenitors independent of hemogenic or capillary sources

To identify the cellular origins that generate kidney lymphatics, we performed genetic lineage tracing using a panel of endothelial-targeted Cre drivers, combined with wholemount immunolabelling for tdTomato and molecular markers of lymphatics. Solvent-based optical clearing and 3D confocal microscopy^48^ were used to visualise lineage contributions at embryonic day (E)15.5, when kidney lymphatic expansion during development is most rapid^29^. We first employed a constitutive *Tie2-Cre* line^49^, which marks venous and capillary endothelium and their progeny. Prior lineage tracing studies demonstrate that *Tie2-Cre* labels LECs in multiple organs^20,33,34,36^, capturing both venous contributions^20^ and an angioblast population that gives rise to systemic LECs^36^. However, *Tie2^+^* lineage labelling of kidney lymphatics has not been explored. In E15.5 *Tie2-Cre*;*Rosa26*^tdTomato/+^ embryos, tdTomato labelled blood vasculature in the metanephric kidney, the direct precursor to the adult organ (**Figure S1a,b**). By co-labelling E15.5 *Tie2-Cre*;*Rosa26*^tdTomato/+^ kidneys with PROX1, PDPN and tdTomato and applying 3D imaging, we manually quantified *Tie2*^+^ progenitor contributions from confocal image stacks of embryonic kidney (**Figure 1a,b**). From 761 LECs examined within five kidneys across two embryonic litters, 643 (86.7 ± 1.8%) kidney LECs were tdTomato⁺, whereas 118 (13.3 ± 1.8%) LECs remained unlabelled.

**Figure 1.**
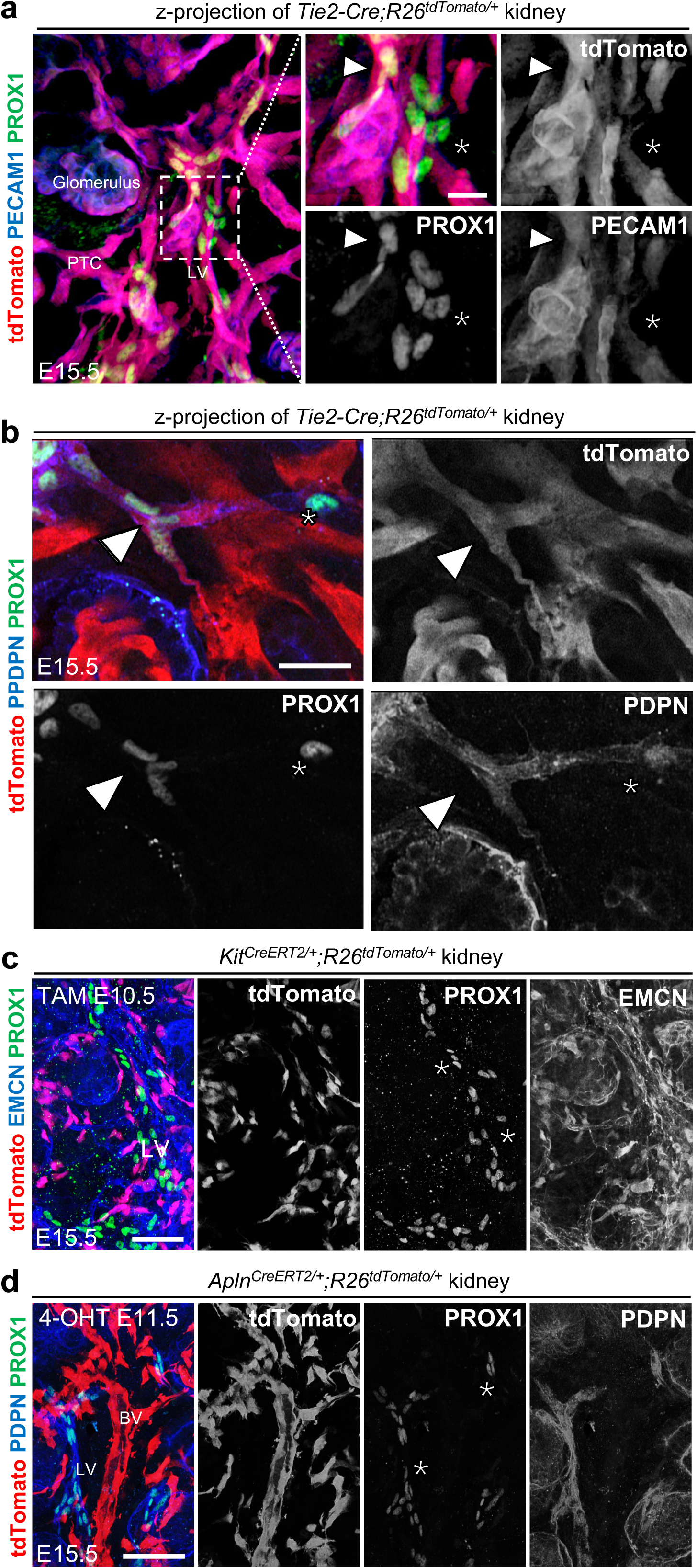
Endothelial progenitor contributions to developing kidney lymphatics. (**a**) Left: Low magnification 3D reconstruction of E15.5 *Tie2-Cre;Rosa26*^tdTomato/+^ kidneys (*n* = 3, two litters), immunolabelled for PROX1, PECAM1 and tdTomato. tdTomato⁺ labelling is observed in the glomerulus, peritubular capillaries (PTC) and lymphatic vessels (LV). Scale bar: 30 μm. Right: Magnified view of dashed box showing tdTomato⁺ LECs (arrowhead) and unlabelled LECs (asterisk). Scale bar: 10 μm. (**b**) Confocal z-section from E15.5 *Tie2-Cre;Rosa26*^tdTomato/+^ kidneys (*n* = 2), labelled for PROX1, PDPN and tdTomato. Some LECs are tdTomato⁺ (arrowhead), but others are unlabelled (asterisk). Scale bar: 20 μm. (**c**) z-projection of E15.5 *Kit*^CreERT2/+^;*Rosa26*^tdTomato/+^ kidneys (*n* = 5, two litters), labelled for PROX1, EMCN and tdTomato following E10.5 tamoxifen. No tdTomato⁺ LECs detected in the cortex. Scale: 50 μm. (**d**) z-projection of E15.5 *Apln*^CreERT2/+^;*Rosa26*^tdTomato/+^ kidneys (*n* = 4), labelled for PROX1, PDPN and tdTomato after E11.5 4-OHT. Robust labelling of blood vessels (BV) observed, but PROX1⁺ LECs remain unlabelled. Scale bar: 50 μm.

*Tie2-Cre* labels multiple vascular beds with potential for lymphatic differentiation. For example, the yolk sac contains hemogenic endothelium with competency for LEC differentiation^34^. Using LYVE1 to demarcate hemogenic endothelial cells^50^, we found LYVE1^+^ EMCN^+^ endothelia were labelled in E11.5 *Tie2-Cre*;*Rosa26*^tdTomato/+^ yolk sac (**Figure S1c**). We therefore used a tamoxifen inducible CreERT2 driver targeting specific endothelial lineages previously shown to contribute to lymphatics. To capture hemogenic endothelium-derived LECs, we used *Kit*^CreERT2/+^ which, when activated at E10.5, labels LECs within the mouse mesentery^28,51^. Using this approach, rare tdTomato⁺ EMCN⁺ blood endothelial cells were observed in the E15.5 kidney (**Figure S2a,b**)^52^ . However, LECs were not tdTomato^+^ (**Figure 1c**, *n* = 3 kidneys from across 2 litters). Tissue-resident blood capillary endothelium has been shown to contribute to lymphatics in the dermis^27^, and we used *Apln*^CreERT2^ to label this potential source^24,53^. 4-hydroxytamoxifen (4-OHT) was administered to *Apln*^CreERT2/+^;*Rosa26*^tdTomato/+^ mice at E11.5, a timepoint that labels the kidney vasculature upon its entry into the metanephric mesenchyme^54^ but precedes the arrival of the lymphatic plexus^29^ . Although this approach led to robust labelling of the EMCN⁺ vasculature (**Figure S3a,b**), kidney LECs did not express tdTomato (**Figure 1d**, *n* = 4 kidneys from across 2 litters).

Together, these findings demonstrate that the majority of kidney lymphatics arise from a *Tie2^+^* endothelial progenitor lineage that is independent of hemogenic endothelium or kidney blood capillaries.

### *Osr1^+^* progenitors account for the remaining proportion of kidney lymphatics and are distinct from other putative lymphatic origins

The mouse metanephric kidney forms around E10.5 and is composed of posterior intermediate mesoderm-derived metanephric mesenchyme and anterior intermediate mesoderm-derived ureteric bud. The posterior intermediate mesoderm gives rise to specialised kidney epithelia, stroma, perivascular cells and blood endothelium^46^, but its putative contribution to LECs has not been explored.

We first ascertained if LEC progenitors exist within intermediate mesoderm. To do this, we interrogated a published single-nucleus RNA-sequencing atlas of mouse embryogenic development^55^, analysing a 95,226 cell subset of the atlas containing intermediate mesoderm and its derivatives from E8.0 until birth (**Figure S4a,b**). The transcription factor *Osr1*, an established marker of intermediate mesoderm essential for kidney development^56–58^, was enriched within cells annotated as posterior intermediate mesoderm or metanephric mesenchyme (**Figure S4c**). Within this population, we identified cells co-expressing *Prox1* and *Vegfr3* (**Figure S4d,e**), two early regulators of LEC identity that are expressed by the first lymphatic-fated cells emerging from the cardinal vein in mouse embryos^19,59^.

We then tested whether kidney LECs originate from the intermediate mesoderm by performing genetic lineage tracing, utilising mice harbouring a tamoxifen-dependent CreERT2 allele that is knocked-in to the endogenous *Osr1* locus^46^. Importantly, OSR1 is not expressed by LECs during embryonic development^60^, avoiding unexpected labelling of LECs. Tamoxifen administration of *Osr1*^eGFP-CreERT2/+^;*Rosa26*^tdTomato/+^ at E9.5 (**Figure S5a**) resulted in widespread tdTomato expression in the metanephric kidney, including PDPN⁺ glomerular podocytes (**Figure S5b**) and PBX1⁺ stromal cells (**Figure S5c**), confirming broad labelling of *Osr1⁺* posterior intermediate mesoderm^46^. No recombination occurred in Cre-negative controls, although scarce tdTomato⁺ cells were observed following vehicle-only injection (**Figure S5a**), demonstrating the importance of including both controls in lineage labelling studies. Following induction at E8.5 or E9.5 to label posterior intermediate mesoderm-derived cells, we consistently detected tdTomato⁺ traced cells co-expressing the LEC markers PROX1 and PDPN in the kidney lymphatic plexus at E15.5 (**Figure 2a,b**; staining present in 5/5 kidneys examined from three independent litters). No co-labelled cells were detected after tamoxifen injection at E10.5 (**Figure 2b**, findings from 3/3 kidneys from two litters for both E9.5 and E10.5). Quantification demonstrated a temporal restriction of tdTomato⁺ labelled-LECs by *Osr1⁺* progenitors. Following E8.5 tamoxifen induction, 162 out of 1,007 (15.6 ± 2.6%) were tdTomato⁺, dropping to 77 out of 1,432 (5.4 ± 0.7%) of tdTomato⁺ after E9.5 induction (mean difference = 10.1%, 95% CI = 6.2-14.1, *p* = 0.0002; **Figure 2c**). The contribution after tamoxifen induction at E8.5 closely matched the proportion of LECs unlabelled by *Tie2*⁺ endothelial progenitors (**Figure 2d**). No tdTomato⁺ labelled-LECs were detected in the kidney after tamoxifen induction at E7.5, capturing anterior intermediate mesoderm-derived cells (data not shown).

**Figure 2.**
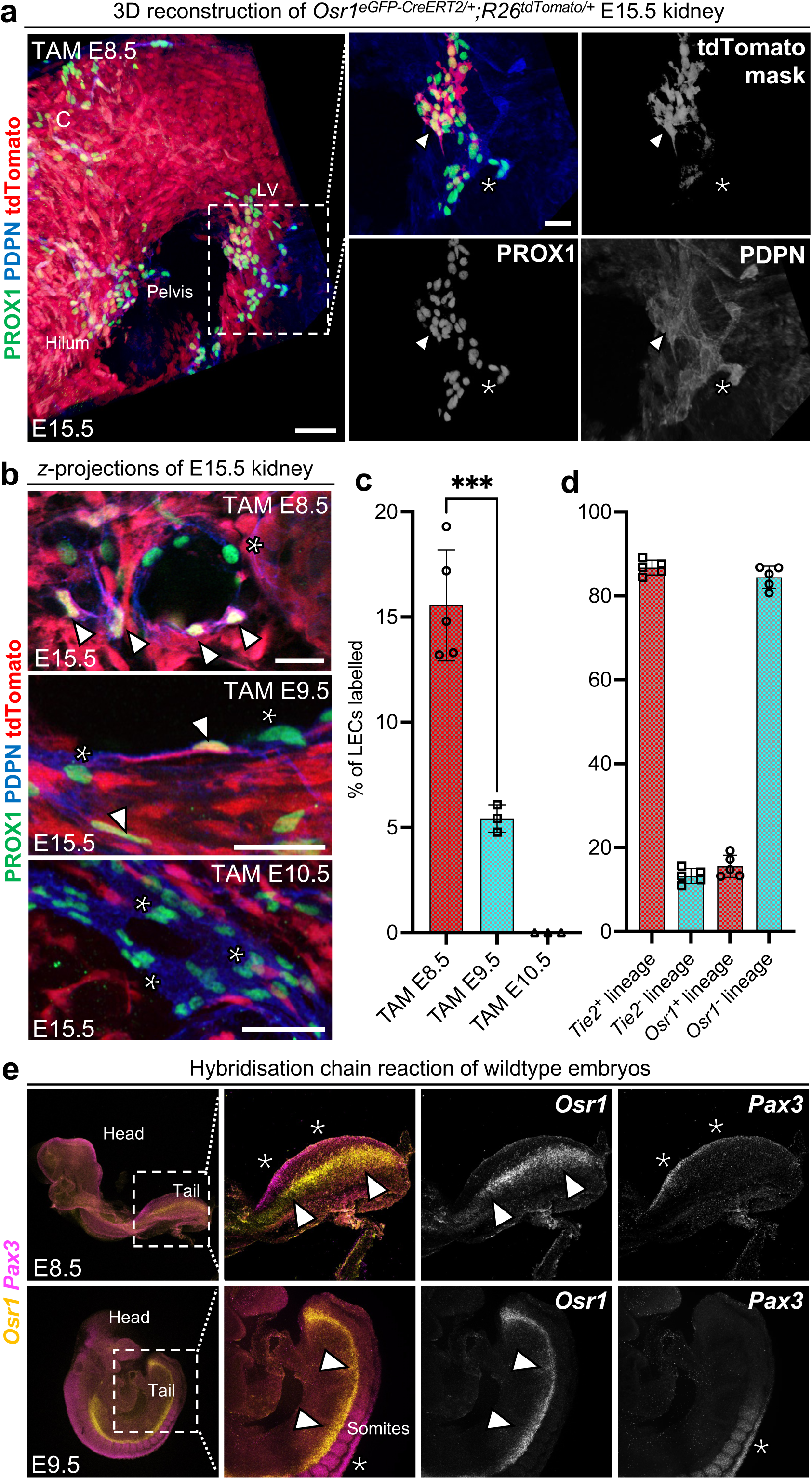
*Osr1*^+^ progenitors contribute to the developing kidney lymphatic plexus. (**a**) Left: Low magnification 3D reconstruction of E15.5 *Osr1*^eGFP-CreERT2/+^;*Rosa26*^tdTomato/+^ kidneys (n = 5, two litters), immunolabelled for PROX1, PDPN and tdTomato after tamoxifen induction at E8.5. Lymphatic vessels (LV) are visible in the hilum and cortex (C). Scale bar: 50 μm. Right: High-resolution reconstruction of boxed region showing tdTomato⁺ and tdTomato⁻ LECs within the PDPN^+^ network (arrowhead and asterisk, respectively). Scale bar: 20 μm. (**b**) Representative z-sections of kidneys following tamoxifen at E8.5, E9.5, or E10.5. Arrowheads indicate tdTomato⁺ LECs. No labelling was seen after E10.5. n ≥ 3 kidneys per timepoint from ≥2 litters. Scale bars: 30 μm. (**c**) Quantification of tdTomato⁺ LECs shows a significant decline in lineage contribution over time (ANOVA F = 68.7, p < 0.0001). E8.5 vs E9.5: mean difference = 10.1%, p = 0.0002. Each point represents one kidney from a separate embryo. \**p* < 0.0332, \*\**p* < 0.0021, \*\*\**p* < 0.0002, \*\*\*\**p* < 0.0001. (**d**) Comparison of proportions of lineage labelled and unlabelled kidney LECs within *Osr1*^eGFP-CreERT2/+^;*Rosa26*^tdTomato/+^ kidneys (labelled: 15.6 ± 2.6% LECs, unlabelled: 84.4 ± 2.6% LECs) compared with *Tie2-Cre;Rosa26*^tdTomato/+^ kidneys (labelled: 86.7 ± 1.8% LECs, unlabelled: 13.3 ± 1.8% LECs). Each data point represents an individual kidney from a separate embryo. (**e**) Wholemount HCR showing *Osr1* and *Pax3* expression at E8.5 (top) and E9.5 (bottom). *Osr1*⁺ intermediate mesoderm (arrowheads) and *Pax3*⁺ somites (asterisks) are spatially distinct. Scale bars: 200 μm.

We next determined whether *Osr1⁺* progenitor-derived kidney LEC originate from non-intermediate mesoderm sources that are known to contribute to lymphatics in other organs. First, we asked whether *Osr1* is expressed in paraxial mesoderm, a major source of systemic LECs^35,36^. Wholemount in situ hybridisation chain reaction (HCR) of wildtype embryos for *Pax3*, a marker of paraxial mesoderm, and *Osr1* at E8.5 and E9.5 (*n =* 5 embryos per timepoint obtained from two litters) revealed spatially distinct expression domains. Between E8.5 and 9.5, *Pax3* expression was detected, as expected, in paraxial mesoderm-derived somites (**Figure 2e**), adjacent to but not overlapping with *Osr1*^+^ cells, especially in the posterior part of the embryo where the *Osr1*^+^ intermediate mesoderm is located. Given the hemogenic endothelial origin of LECs in the heart^34^ and mesentery^28^, we also examined yolk sacs from *Osr1*^eGFP-CreERT2/+^;*Rosa26*^tdTomato/+^ embryos after tamoxifen induction E8.5. The E11.5 yolk sac LYVE1^+^ haemogenic endothelium, did not contain tdTomato⁺ cells (**Figure S5d**).

These results demonstrate that *Osr1⁺* progenitor cells contribute to kidney LECs within a defined developmental window. This lineage reflects a putative intermediate mesoderm origin and accounts for the proportion of LECs unlabelled by the *Tie2^+^*endothelial lineage.

### The *Osr1^+^* lineage does not contribute to lymphatic endothelial cells in the mesentery, heart or dermis

To assess whether *Osr1⁺* progenitors contribute to organ LECs beyond the kidney, we examined organs within which novel LEC progenitors have been identified, namely the heart^24,25,33,34^, dermis^26,27,61^ and mesentery^28^. We profiled organs from the same four E15.5 *Osr1*^eGFP-CreERT2/+^;*Rosa26*^tdTomato/+^ embryos in which we had observed tdTomato⁺ LECs in the kidney.

Wholemount immunolabelling and optical clearing confirmed the presence of PROX1⁺ PDPN⁺ lymphatic networks in each organ. In the heart, PROX1⁺ lymphatics were evident proximal to the atria and extending into the ventricles (**Figure 3a**)^51^, but no tdTomato⁺ LECs were detected in any of the four hearts examined. In the dorsal skin, tdTomato⁺ perivascular cells were found adjacent to dermal lymphatics, as expected^46^, but PROX1⁺ PDPN⁺ LECs lacked tdTomato expression (**Figure 3b**). In the mesentery and adjacent intestinal tracts, PROX1⁺ lymphatic vessels extended into the parenchyma, but no *Osr1^+^* lineage labelled LECs were detected, although rare tdTomato⁺ cells lacking lymphatic markers were seen near vessels (**Figure 3c**).

**Figure 3.**
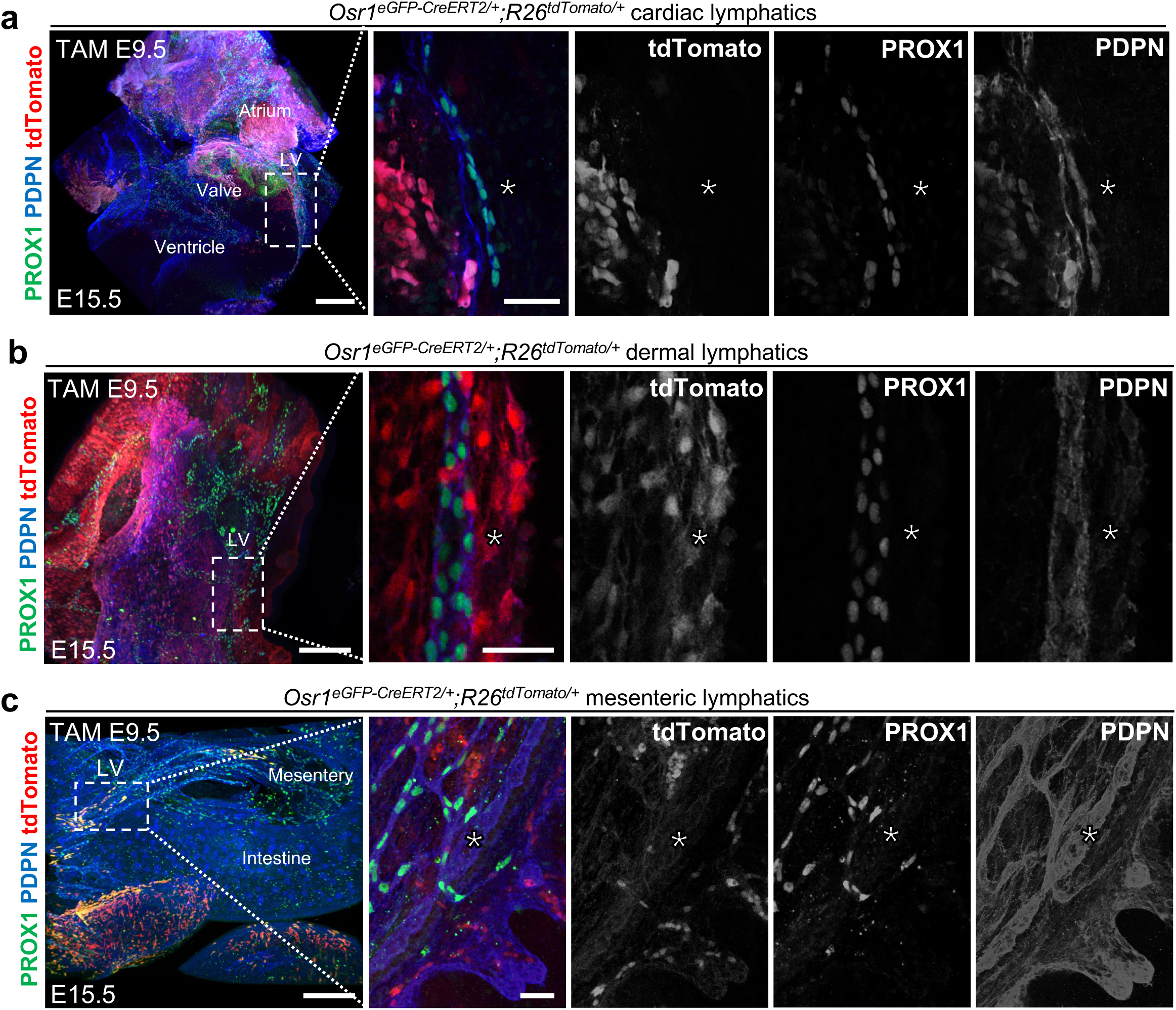
The *Osr1^+^* lineage does not contribute to lymphatics in the heart, skin or mesentery. (**a**) Left: low magnification z-projection of E15.5 *Osr1*^eGFP-CreERT2/+^;*Rosa26*^tdTomato/+^ heart, pooled from two litters and immunolabelled for PROX1, PDPN and tdTomato. Lymphatic vessels (LV) extend from the atria to ventricles, with PROX1⁺ valve cells visible. Scale bar: 200 μm. Right: high magnification of boxed region from (a), showing absence of tdTomato labelling in a cardiac lymphatic vessel (asterisk). Scale bar: 30 μm. (**b**) Left: Low magnification z-projection of dorsal skin from E15.5 embryos, showing dermal lymphatics (LV) labelled for PROX1, PDPN and tdTomato. Scale bar: 200 μm. Right: high magnification of boxed region from (c), confirming no tdTomato⁺ LECs (asterisk). Scale bar: 30 μm. (**c**) Left: low magnification z-projection of E15.5 mesentery and adjacent intestine (Int), showing lymphatic vessels extending from mesentery (M). Scale bar: 200 μm. Right: high magnification of boxed region from (e), showing absence of tdTomato expression in PROX1⁺ PDPN⁺ lymphatic vessels (asterisk). Scale bar: 30 μm. Tamoxifen was administered at E9.5. For each tissue, a minimum of two samples from independent litters were analysed, with all imaging representative of at least three fields of view per sample.

These results demonstrate that the *Osr1⁺* lineage does not contribute to LECs in the heart, skin, or mesentery, in contrast with its specific contribution to the kidney. This organ-restricted pattern supports the idea that *Osr1⁺* lineage LECs arise from intermediate mesoderm rather than a shared or systemic progenitor source.

### *Osr1^+^* lineage lymphatics arise independently of stromal or nephron lineages within the kidney

During kidney development, the *Osr1^+^* posterior intermediate mesoderm segregates into epithelial and stromal compartments. Of these two lineages, stromal progenitors expressing FOXD1^62–64^ generate vascular smooth muscle, pericytes and interstitial stromal cells, with a subset of progenitors expressing TBX18^65,66^ forming differentiated stromal cell types within the kidney hilum. We therefore examined whether, downstream of posterior intermediate mesoderm, kidney lymphatics derive from the *Foxd1⁺* lineage of stromal progenitors. Using a constitutive *Foxd1*^eGFP-Cre/+^ line, we found tdTomato⁺ labelling of PBX1⁺ stromal cells in *Foxd1*^eGFP-Cre/+^;*Rosa26*^tdTomato/+^ kidneys at E15.5 (**Figure S6a,b**). We also found that a subset of kidney LECs (**Figure 4a**) were tdTomato^+^ at this timepoint.

**Figure 4.**
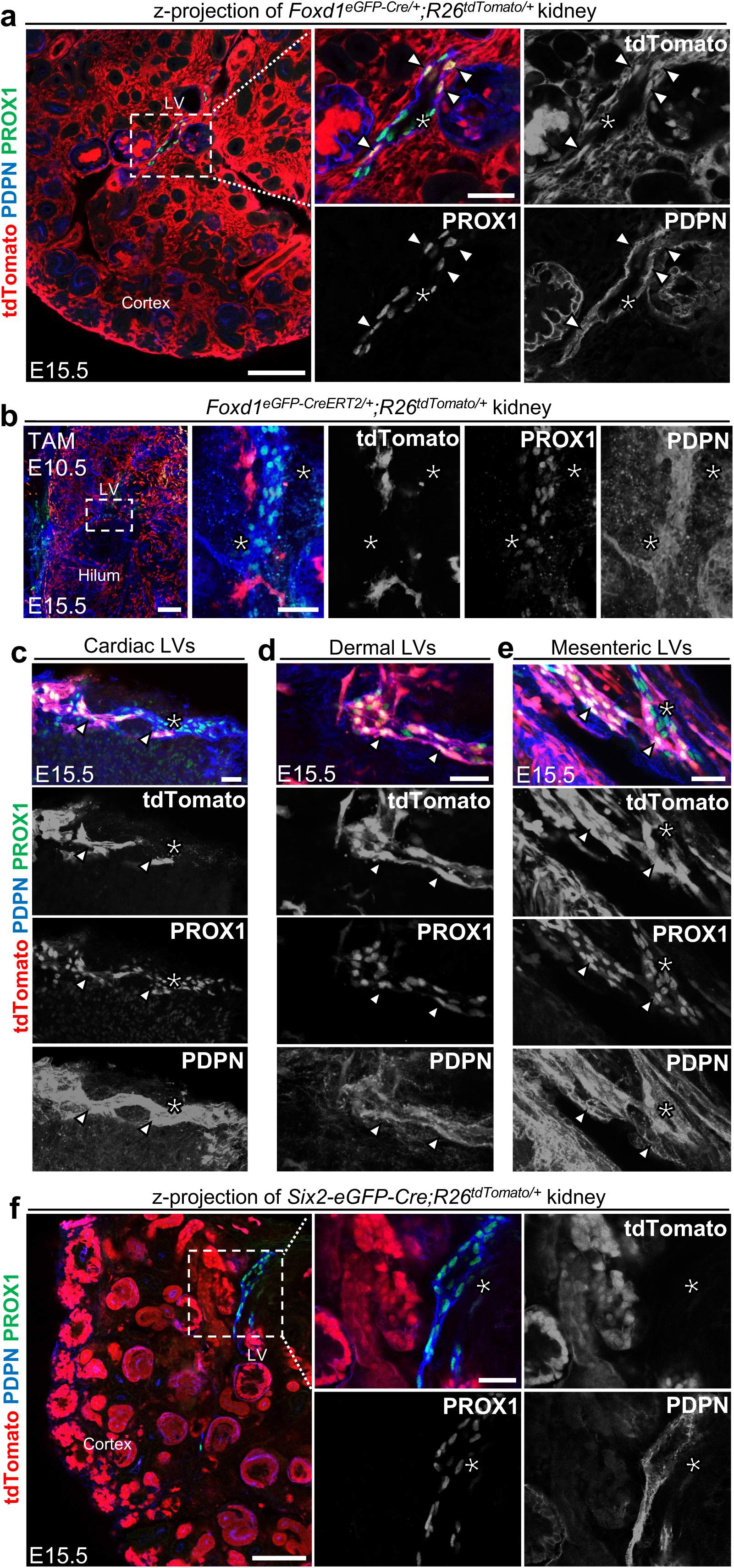
Examining nephron and stromal lineage contributions to kidney lymphatics. (**a**) Left: low magnification z-section of E15.5 *Foxd1*^eGFP-Cre/+^;*Rosa26*^tdTomato/+^ kidneys (n = 5; two litters) immunolabelled for PROX1, PDPN and tdTomato. Lymphatics (LV) were seen en route to the cortex. Scale bar: 100 μm. Right: high-resolution image of boxed region reveals tdTomato⁺ PROX1⁺ PDPN⁺ LECs (arrowheads), interspersed with unlabelled cells (asterisks), indicating *Foxd1^+^* lineage contribution. Scale bar: 30 μm. (**b**) Left: representative low-resolution z-projection of E15.5 *Foxd1*^eGFP-CreERT2/+^;*Rosa26*^tdTomato/+^ kidneys (*n* = 4), immunolabelled for PROX1, PDPN and tdTomato. Lymphatic vessels (LV) are shown in the hilum adjacent to the pelvis. Scale bar: 100 μm. Right: magnified dashed box region from left panel, showing no overlap between tdTomato, PROX1 and PDPN in lymphatic vessels (asterisk). Scale bar: 30 μm. (**c-e**) High magnification z-projections of E15.5 *Foxd1*^eGFP-Cre/+^;*Rosa26*^tdTomato/+^ tissues: heart (**c**), skin (**d**) and mesentery (**e**), pooled from two litters (*n* = 2 organs per group), immunolabelled for PROX1, PDPN and tdTomato. tdTomato⁺ LECs (arrowheads) were observed in each organ, coexisting with non-lineage-labelled LECs (asterisks), reflecting *Foxd1*⁺ lineage contribution beyond the kidney. Scale bar: 30 μm. (**f**) Left: low magnification z-section of E15.5 *Six2-eGFP-Cre;Rosa26*^tdTomato/+^ kidneys (*n* = 4; two litters), immunolabelled for PROX1, PDPN and tdTomato. Lymphatic vessels (LV) were visible extending into the cortex. Scale bar: 100 μm. Right: high-resolution image of boxed region, showing no tdTomato expression in PROX1⁺ PDPN⁺ LECs (asterisk), indicating absence of contribution from the *Six2*⁺ nephron progenitor lineage. Scale bar: 30 μm.

To ascertain whether these *Foxd1⁺* lineage lymphatics represented a kidney stromal progenitor contribution, we sought to replicate our findings using inducible *Foxd1*^eGFP-CreERT2^ mice. This CreERT2, when labelled with tamoxifen at E10.5, captures both cortical and medullary kidney stromal progenitors^63^. In contrast to our observations with the constitutive Cre line, PROX1⁺ PDPN⁺ LECs were not labelled with the inducible approach (**Figure 4b**, three kidneys from two litters). We then utilised the inducible *Tbx18*^CreERT2/+^ line^67^ that, upon tamoxifen induction, marks FOXD1⁺ stromal progenitors in the kidney hilum^65,66^, an early site of lymphatic colonisation^29^. Induction at E10.5 caused tdTomato⁺ labelling of perivascular stroma and smooth muscle at the kidney hilum (**Figure S7a**), but tdTomato⁺ LECs were not detected (**Figure S7b**, 3/3 kidneys), excluding stromal lineage contributions to kidney LECs.

The absence of kidney lymphatic labelling using these approaches led us to hypothesise that *Foxd1⁺* lineage LECs arise from a cellular source that is distinct from *Osr1^+^* intermediate mesoderm. Evidence for *Foxd1* expression in the paraxial mesoderm in non-mammalian species^68,69^ prompted us to explore its expression in mouse embryos. Wholemount HCR of wildtype embryos showed that *Foxd1* was expressed from E8.0-8.5 (not shown) and at E9.5 within paraxial mesoderm-derived somites and was spatially segregated from the *Osr1*^+^ intermediate mesoderm field (**Figure S8,** *n =* 3 embryos from two litters). In line these findings, across three *Foxd1*^eGFP-Cre/+^;*Rosa26*^tdTomato/+^ embryos, we consistently detected tdTomato⁺ LECs in the heart (**Figure 4c**), dermis (**Figure 4d**) and mesentery (**Figure 4e**), organs in which paraxial mesoderm is known to contribute to lymphatics^35,36^ and differing from our experiments demonstrating a kidney-restricted contribution of the *Osr1*^+^ lineage. Thus, *Foxd1⁺* lineage LECs are most likely to arise from paraxial mesoderm, rather than kidney stroma downstream of the *Osr1^+^* lineage.

The second major lineage within the *Osr1^+^* posterior intermediate mesoderm is demarcated by SIX2 expression, serving as a progenitor source for nephron epithelia and glomerular podocytes^70^. To determine whether the nephron lineage contributes to kidney lymphatics, we utilised a constitutively active *Six2-eGFP-Cre*. In *Six2-eGFP-Cre*;*Rosa26*^tdTomato/+^ embryos at E15.5, tdTomato robustly labelled ECAD⁺ kidney epithelia (**Figure S9a,b**), but not PROX1⁺ PDPN⁺ LECs (**Figure 4f**) across four kidneys examined from two litters.

Taken together, these findings indicate that, akin to the blood vasculature within the kidney^46^, the small proportion of kidney lymphatics that derive from the *Osr1*^+^ progenitor pool do so directly, independent of *Foxd1^+^* stromal or *Six2^+^* nephron lineages.

### Kidney lymphvasculogenesis involves contributions from both *Tie2⁺* and *Osr1⁺* progenitor lineages

Previously, we identified anatomically distinct clusters of LECs in the mouse and human developing kidney, suggestive of *de novo* formation of lymphatics *via* lymphvasculogenesis^14,22^. To investigate the lineage composition of these clusters, we first assessed the contribution of the *Tie2⁺* endothelial lineage, given its broad contribution to kidney LECs. From nine non-overlapping 3D imaging volumes of *Tie2-Cre*;*Rosa26*^tdTomato/+^ kidneys at E15.5, 24 out of 37 (64.0 ± 9.1%) PROX1⁺ PDPN⁺ LECs across thirteen lymphatic clusters (from three embryos) were tdTomato⁺ (**Figure 5a**), indicating that the *Tie2⁺* endothelial lineage contributes substantially to *de novo* lymphatic formation.

**Figure 5.**
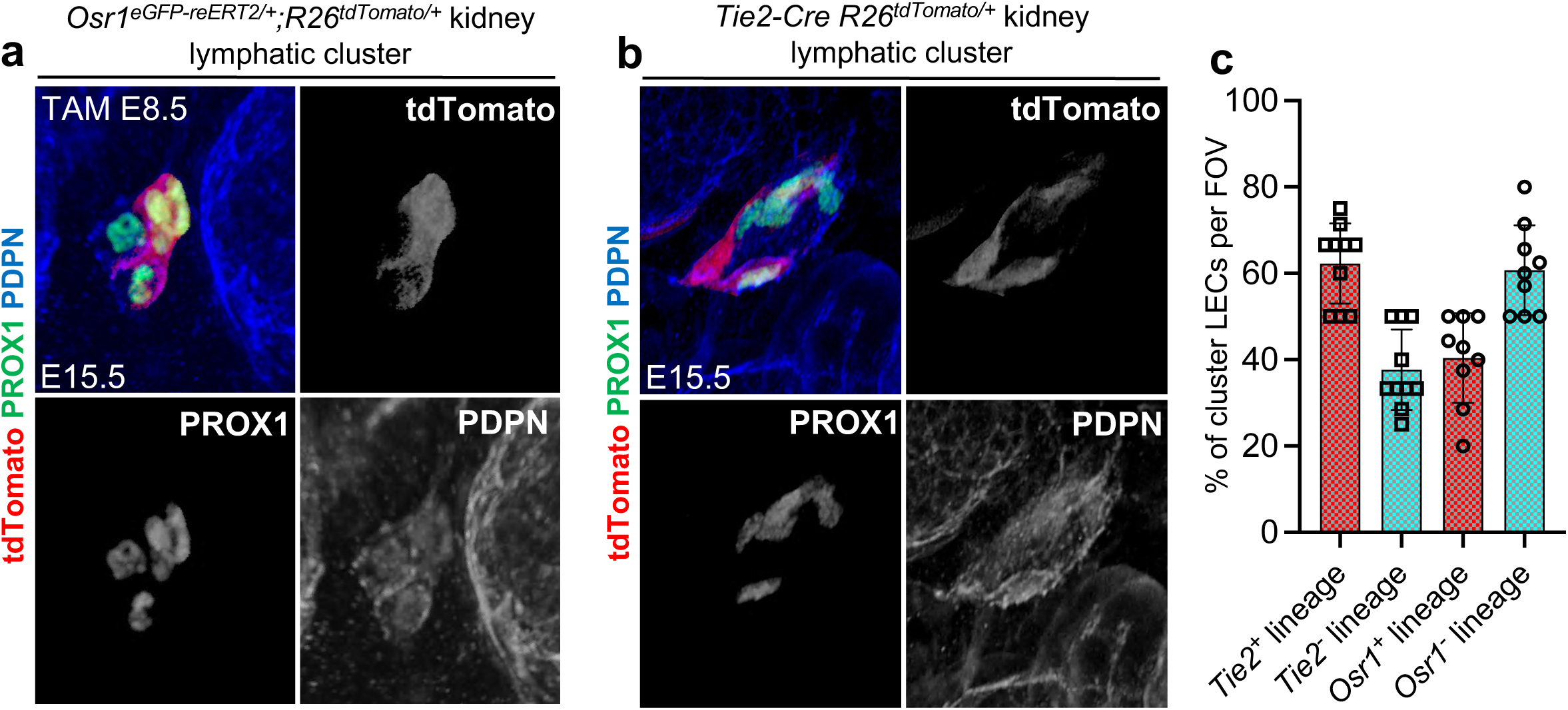
*Osr1*^+^ and *Tie2^+^* progenitors contribute to kidney lymphvasculogenesis. (**a**) 3D reconstruction of 13 lymphatic clusters from *Tie2-Cre;Rosa26*^tdTomato/+^ kidneys (E15.5), pooled from two litters and labelled for PROX1, PDPN and tdTomato. tdTomato⁺ cluster LECs (arrowheads) co-express PROX1 and PDPN, whereas others remained unlabelled by tdTomato (asterisks). Representative of nine non-overlapping 3D imaging volumes. Scale bar: 10 μm. (**b**) Representative 3D reconstruction of 16 lymphatic clusters from *Osr1*^eGFP-CreERT2/+^;*Rosa26*^tdTomato/+^ kidneys (E15.5), pooled from two litters and immunolabelled for PROX1, PDPN and tdTomato. Tamoxifen was administered at E8.5. tdTomato⁺ LECs (arrowheads) are intermixed with non-labelled LECs (asterisks) within clusters. Representative of eight non-overlapping 3D imaging volumes. Scale bar: 10 μm. (**c**) Quantification of lineage-labelled or unlabelled LECs within lymphatic clusters at E15.5. In *Osr1*^eGFP-CreERT2/+^;*Rosa26*^tdTomato/+^ kidneys, 36.2 ± 10.4% of cluster LECs were lineage labelled and 63.8 ± 10.4% were unlabelled. Each data point represents an individual field of view in which clusters were assessed.

We next examined the contribution of *Osr1⁺* intermediate mesoderm to lymphvasculogenesis in the kidney. Using *Osr1*^eGFP-CreERT2/+^;*Rosa26*^tdTomato/+^ kidneys after tamoxifen induction at E8.5, and from eight non-overlapping imaging volumes, we found that 24 out of 59 (36.2 ± 10.4%) PROX1⁺ PDPN⁺ LECs within sixteen clusters (from three embryos) were tdTomato⁺ (**Figure 5b**). Additionally, two of the sixteen clusters were exclusively comprised of *Osr1*⁺ lineage-derived cells. Quantification revealed that the proportion of LECs within kidney lymphatic clusters that were labelled by the *Osr1*⁺ lineage was complementary to *Tie2⁺* lineage-negative LECs within these clusters (Figure 5a).

These findings suggest that *de novo* LEC formation in the kidney, *via* lymphvasculogenesis, involves contributions from both *Tie2⁺* endothelial and *Osr1*⁺ intermediate mesodermal progenitors.

## DISCUSSION

Our understanding of lymphatic origins has expanded considerably over the past decade. Classical studies using *Tie2-Cre* mice implicated a predominantly venous origin for LECs^20^, consistent with earlier dye injection experiments in mammalian embryos^15,16^. However, subsequent lineage tracing has revealed surprising diversity in the cellular origins of LECs, with sources including the second heart field^25,33^, hemogenic endothelium^28,34^, tissue-resident blood capillaries^27^ and paraxial mesoderm^35,36^. Here, we expand this framework by identifying two distinct and complementary progenitor sources for kidney lymphatics: a *Tie2⁺* endothelial lineage, which contributes the majority of LECs and is shared by other organs, and an *Osr1⁺* intermediate mesoderm lineage, representing a previously unrecognised, kidney-specific LEC source. Unlike other organs profiled to date, the kidney contains LECs from a progenitor pool not shared with systemic lymphatic beds.

Our study places lymphatic development within the broader framework of lineage hierarchies within the kidney. Using snRNA-seq, we identified *Prox1* and *Vegfr3* expression within subsets of *Osr1⁺* intermediate mesoderm, supporting early lymphatic fate specification within this population. Subsequently, our genetic lineage tracing studies reveal that, in addition to giving rise to nephron epithelium, stroma, and blood vasculature^46^, the posterior intermediate mesoderm also gives rise to ∼15% of kidney LECs. This contribution was found to arise after tamoxifen administration at E8.5. This timing precedes the emergence of the definitive metanephric rudiment at E10.5, the direct precursor to the adult kidney. However, at E10.5, no lineage labelling was identified, in line with the known temporal lineage restriction of *Osr1⁺* cells as kidney development progresses^46^. Importantly, the *Osr1⁺* lineage contribution was exclusive to the kidney. *Osr1⁺-*derived LECs were not detected in the heart, mesentery, or dermis, organs in which alternative mesodermal progenitors such as the second heart field and paraxial mesoderm contribute to lymphatics^25,33,35,36^. As *Osr1* is expressed in non-kidney progenitors, including posterior second heart field^71^ and fibroblast precursors^60,72^, the detection of *Osr1⁺* progenitor-derived lymphatics in the kidney, as opposed to other organs examined, suggests that *Osr1⁺-*derived LECs are unique to the kidney and arise from intermediate mesoderm, rather than shared progenitor populations

To determine whether kidney LECs emerge downstream of known lineages within the intermediate mesoderm, we performed lineage tracing of *Foxd1⁺* stromal and *Six2⁺* nephron progenitor populations. While constitutive *Foxd1*^eGFP-Cre^ labelled some kidney LECs, this labelling was absent in the inducible *Foxd1*^eGFP-CreERT2^ line. Instead, *Foxd1* expression was enriched within *Pax3^+^* paraxial mesoderm–derived somites at early timepoints, and *Foxd1*⁺ LECs were detected in organs within which paraxial mesoderm is known to contribute to lymphatics^35,36^. These data suggest that *Foxd1*^eGFP-Cre^ captures paraxial mesoderm derivatives rather than kidney-specific stroma *per se*. Additionally, *Six2⁺* nephron progenitors did not contribute to kidney LECs.

Taken together, our results suggest that *Osr1⁺* lineage LECs arise independently of canonical stromal or epithelial compartments within the kidney. This mirrors the developmental pattern of renal blood vasculature, which also arises from the *Osr1⁺* ^46^, but not *Foxd1⁺* or *Six2⁺* lineages^64,70^. Our data may also clarify discrepancies in earlier lineage tracing studies of the kidney. While one study report found that *Foxd1*^eGFP-Cre^ labelled CD31⁺ endothelial cells in the kidney^62^, another found no *Foxd1⁺* lineage contribution to renal blood vessels^64^. Given that LECs can express CD31, albeit at lower levels than blood endothelial cells^29^, it is plausible that some of the previously identified CD31⁺ cells were LECs. These findings underscore the importance of temporally controlled lineage tracing to avoid misattributing cellular origins.

A key finding is that both *Tie2⁺* and *Osr1⁺* lineages contribute to lymphatic clusters within the developing kidney, with complementary proportions of lineage labelling. Anatomically discrete clusters of LECs have previously been observed in the developing meninges^23^, heart^24,25^, skin^26,27^, mesentry^28^ and kidney^29^. Inspired by live imaging studies within zebrafish^30–32^, these clusters have been proposed to arise *via* lymphvasculogenesis; *de novo* LEC differentiation from tissue-resident progenitors. An analogous mechanism has been proposed for the blood vasculature of the kidney, which forms both by sprouting^54,73^ and *de novo* differentiation of kidney endothelial cells *in situ*^74–78^. Although once thought to reflect non-venous origins, recent evidence shows that paraxial mesoderm–derived progenitors can contribute both to lymphatic sprouting and to cluster formation^35,36^. Our data extend this concept by showing that the kidney hosts lymphatic clusters composed of both a systemic *Tie2⁺* lineage and a local, kidney-specific *Osr1⁺* lineage. Thus, cluster formation *via* lymphvasculogenesis appears not to depend on a singular lineage identity, but rather represents a shared cellular mechanism across progenitor types.

There are limitations to our study. As with all Cre-based lineage tracing, conclusions are dependent on the specificity and efficiency of the Cre drivers. We attempted to mitigate this using a range of constitutive and inducible lines, but definitive resolution of lineage relationships may require intersectional genetics^79^ or single-cell barcoding approaches^80,81^. Additionally, our analysis is limited to embryonic development. How these lineages persist or remodel in postnatal or diseased kidneys remains unexplored. Finally, the molecular cues that instruct lymphatic fate within the intermediate mesoderm remain to be elucidated.

In conclusion, we demonstrate that kidney lymphatics arise from dual progenitor sources: a broadly conserved *Tie2⁺* endothelial lineage and a kidney-restricted *Osr1⁺* intermediate mesoderm lineage. Our findings support an emerging model in which lymphatic development is governed by organ-specific lineage hierarchies, akin to those shaping blood vasculature. As with blood endothelial cells, LECs may retain developmental memory that influences their specialised functions in health and disease. Understanding how distinct progenitors establish lymphatic identity may ultimately inform strategies for selectively targeting or regenerating kidney lymphatics, which undergo expansion in response to injury and inflammation^37–40^ but are perturbed in contexts such as chronic transplant rejection^45,82^. Our study provides a foundation for investigating how lymphatic identity is encoded during kidney development and how it might be leveraged in regeneration and repair.

---

**Figure S1.**
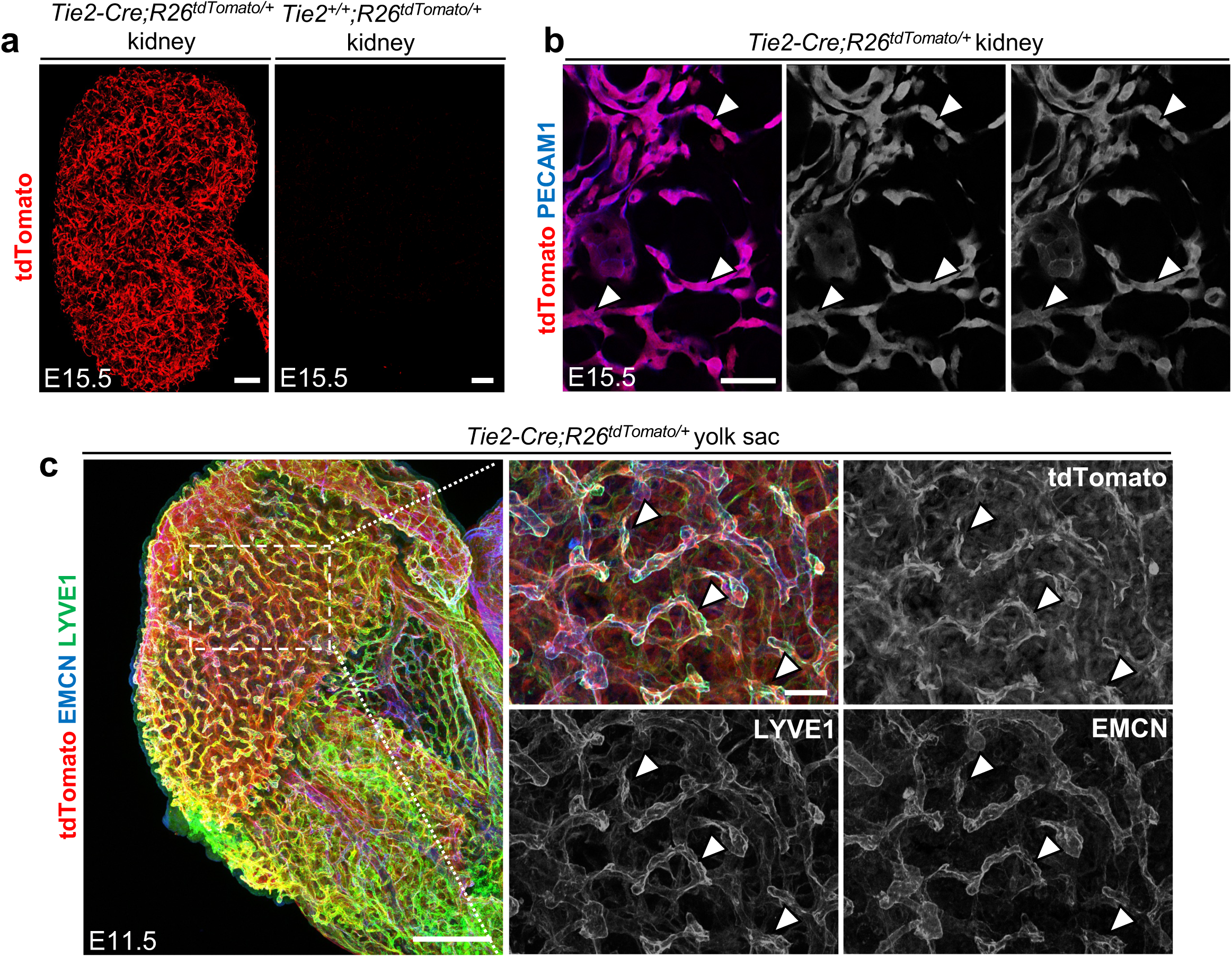
Validation of *Tie2^+^* progenitor lineage labelling in the kidney and yolk sac. (**a**) Representative images from two *Tie2-Cre;Rosa26*^tdTomato/+^ kidneys (left, *n* = 4 fields) and one Cre-negative littermate control (right), immunolabelled for tdTomato. Widespread tdTomato expression is observed in kidney vasculature, with no signal in controls. Scale bar: 100 μm. (**b**) High-resolution z-sections of E15.5 *Tie2-Cre;Rosa26*^tdTomato/+^ kidneys (*n* = 2), immunolabelled for PECAM1 and tdTomato. tdTomato⁺ signal is seen throughout PECAM1⁺ vasculature in the cortex (arrowheads). Scale bar: 30 μm. (**c**) Left: z-projection of E11.5 yolk sacs from *Tie2-Cre;Rosa26*^tdTomato/+^embryos (*n* = 3), labelled for EMCN, LYVE1 and tdTomato. Scale: 200 μm. Right: High-magnification view of boxed region, showing tdTomato⁺ LYVE1⁺ EMCN⁺ cells (arrowheads) in yolk sac vasculature, indicating labelling of hemogenic endothelium by *Tie2-Cre*. Scale bar: 30 μm.

**Figure S2.**
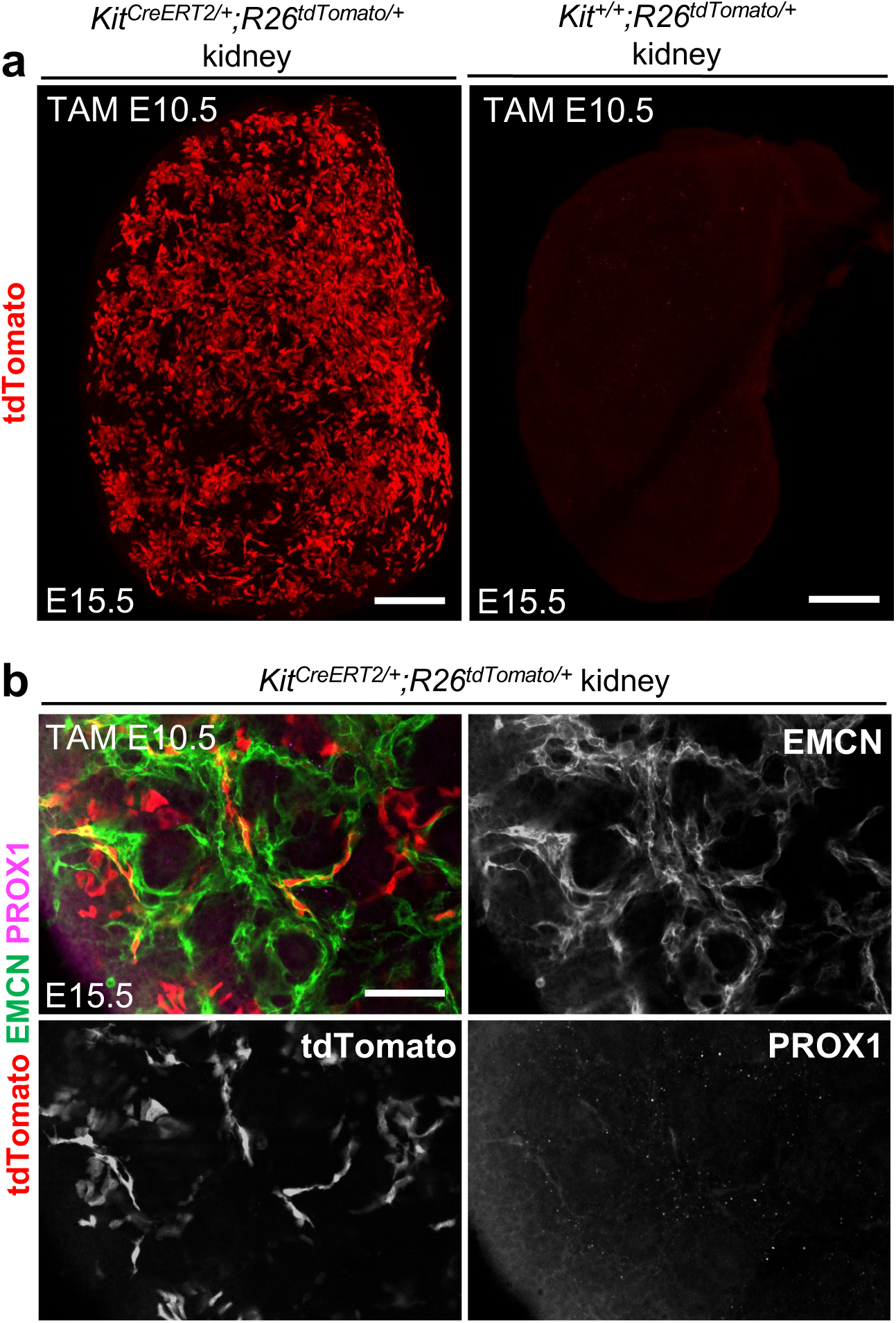
*Kit*^+^ progenitor lineage tracing within in the kidney. (**a**) Representative images of E15.5 *Kit*^CreERT2/+^;*Rosa26*^tdTomato/+^ kidneys (*n* = 2) and a Cre-negative control, immunolabelled for tdTomato. Tamoxifen was administered at E10.5. tdTomato⁺ cells are broadly visible in a vascular pattern; no signal is observed in controls. Scale bar: 100 μm. (**b**) High-resolution confocal z-section from *Kit*^CreERT2/+^;*Rosa26*^tdTomato/+^ kidneys (*n* = 4) immunolabelled for EMCN, PROX1 and tdTomato. tdTomato⁺ cells are occasionally observed within EMCN⁺ blood vessels (arrowheads), but not within PROX1⁺ lymphatics. Scale bar: 50 μm.

**Figure S3.**
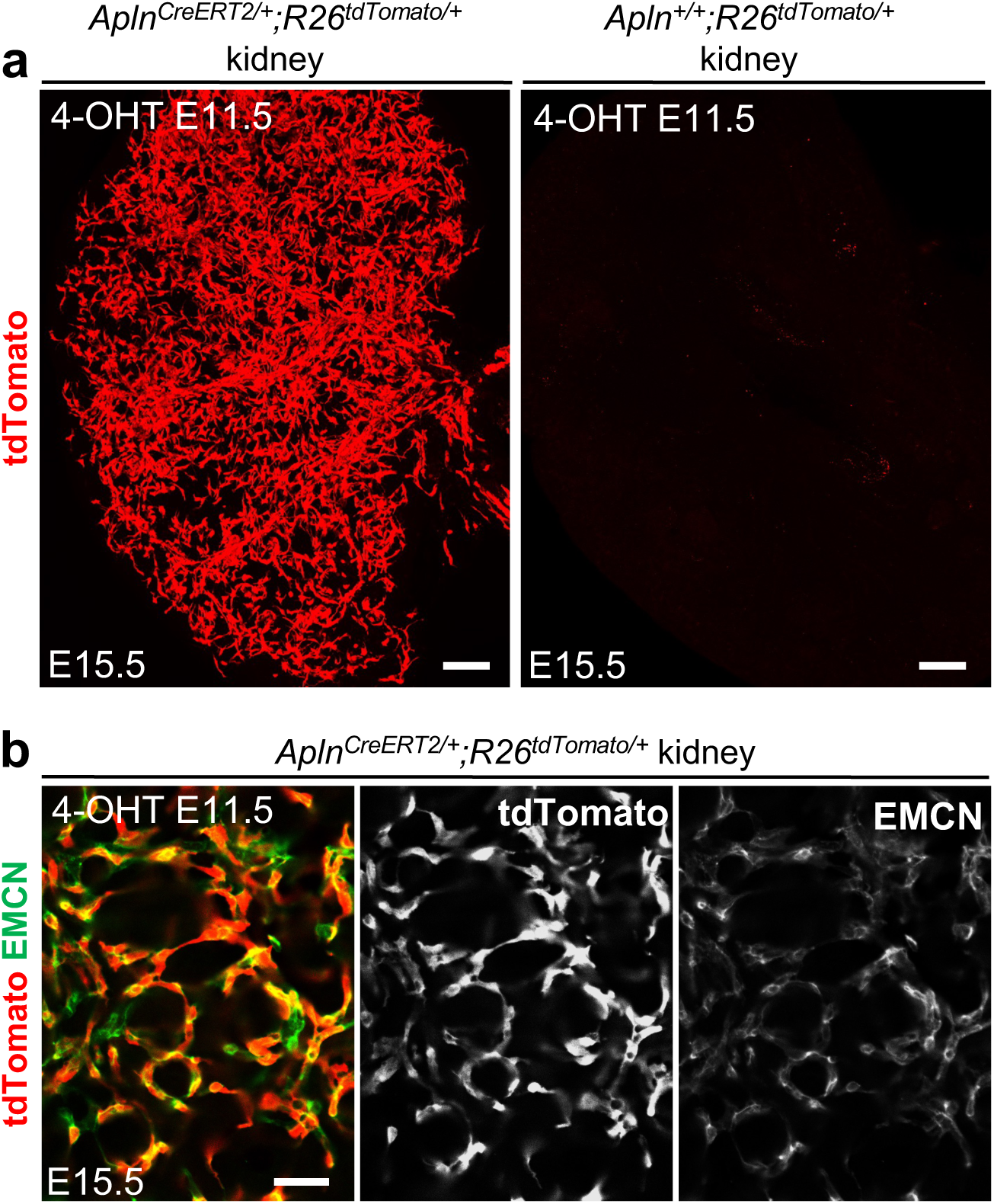
*Apln*^+^ progenitor lineage tracing within in the kidney. (**a**) tdTomato immunolabelling in E15.5 *Apln*^CreERT2/+^;*Rosa26*^tdTomato/+^ kidneys (*n* = 2), following 4-OHT injection at E11.5. Strong vascular tdTomato expression is seen, absent in Cre-negative controls. Scale bar: 100 μm. (**b**) High-resolution z-projection from *Apln*^CreERT2/+^;*Rosa26*^tdTomato/+^ kidneys (*n* = 3), labelled for EMCN and tdTomato. tdTomato⁺ signal colocalises with EMCN⁺ endothelial cells (arrowheads) in cortical vasculature. Scale bar: 30 μm.

**Figure S4.**
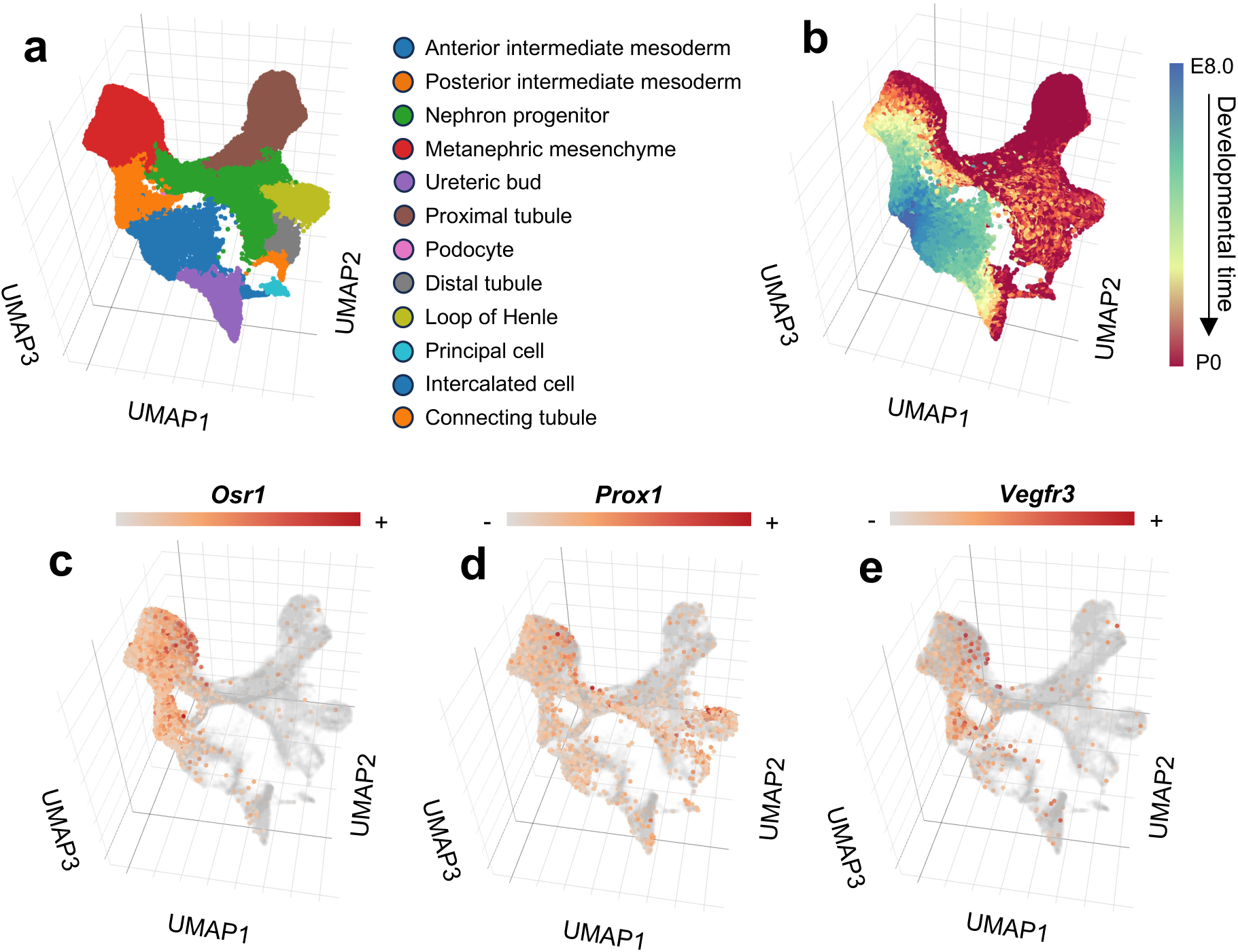
Analysis of single-nucleus RNA-sequencing data from mouse embryos supports lymphatic progenitors within *Osr1*^+^ intermediate mesoderm. (**a-b**) A previously published single-nucleus RNA-sequencing (snRNA-seq) atlas of mouse embryonic development, spanning embryos from E8.0 until the postnatal stage, was interrogated using the online browser: https://omg.gs.washington.edu. 3D uniform manifold approximation and projection (UMAP) of the ‘kidney development’ module within the atlas, coloured by cell type (**a**) and by timepoint (**b**). (**c-e**) Feature plots showing expression of selected transcripts within the dataset. *Osr1* is expressed within posterior intermediate mesoderm and metanephric mesenchyme (**c**), with *Prox1* (**d**) and *Vegfr3* (**e**) expressed in cells within these domains.

**Figure S5.**
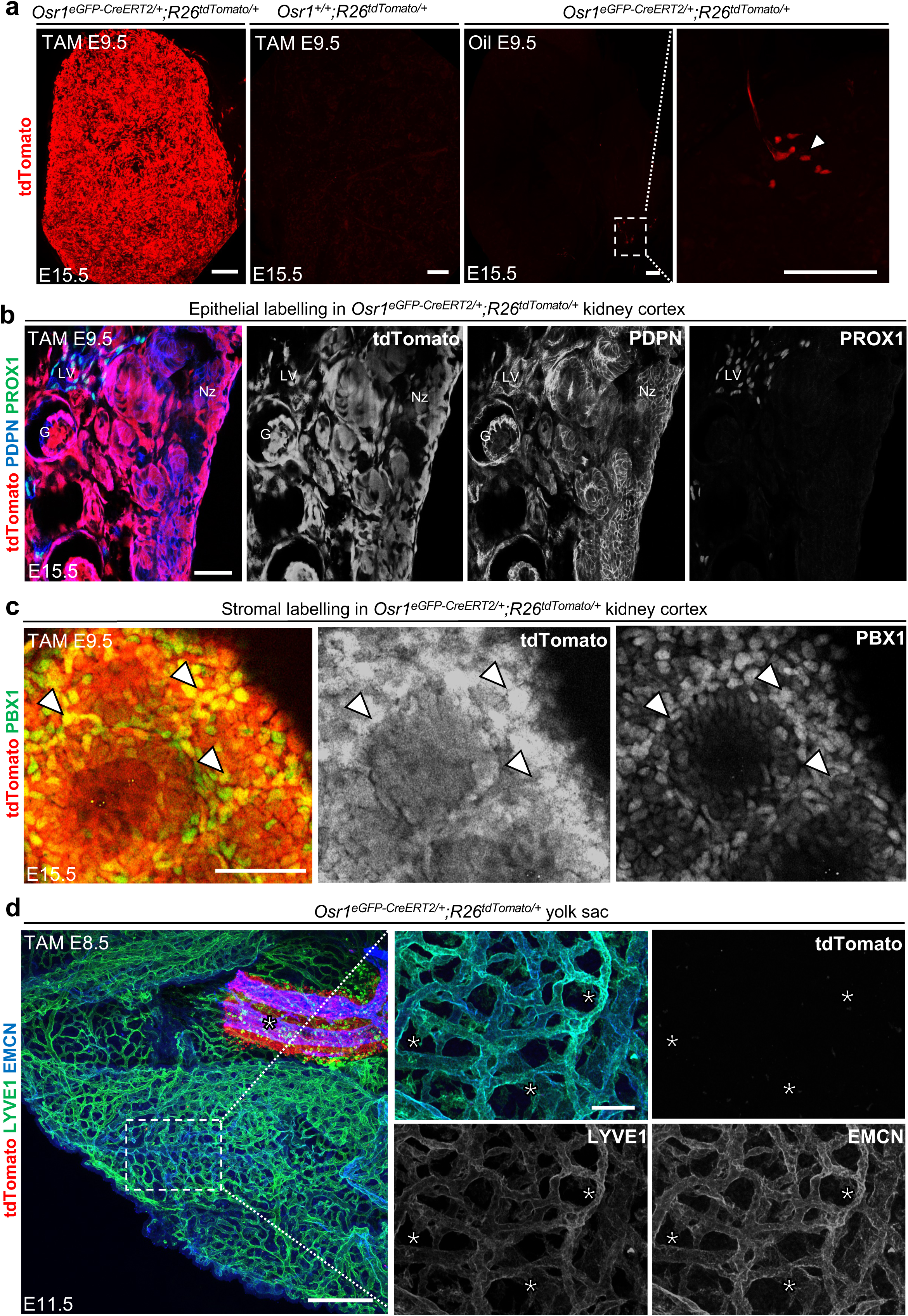
Validation of *Osr1*^+^ progenitor lineage labelling in the kidney. (**a**) Left: representative images of E15.5 *Osr1*^eGFP-CreERT2/+^;*Rosa26*^tdTomato/+^ kidneys (*n* = 4) immunolabelled for tdTomato after tamoxifen induction at E9.5, showing widespread labelling alongside control kidneys (*Osr1*^+/+^;*Rosa26*^tdTomato/+^) lack tdTomato expression. Right: E15.5 kidneys from *Osr1*^eGFP-CreERT2/+^;*Rosa26*^tdTomato/+^ embryos (*n* = 2) injected with oil vehicle at E9.5 exhibit minimal tdTomato expression. Rare tdTomato⁺ cells (arrowheads) indicate low background leakiness of the CreERT2 allele. Magnified view of boxed region is shown. Scale bars: 100 μm. (**b**) High-resolution z-section showing tdTomato⁺ labelling in PROX1⁺ PDPN⁺ epithelia of *Osr1*^eGFP-CreERT2/+^;*Rosa26*^tdTomato/+^ kidneys at E15.5 (*n* = 2), following E9.5 tamoxifen. Signal is seen in glomerular (G) podocytes and the nephrogenic zone (Nz), alongside tdTomato⁺ LECs in lymphatic vessels (LV). Images representative of three fields per kidney. Scale bar: 50 μm. (**c**) High-resolution z-section of E15.5 *Osr1*^eGFP-CreERT2/+^;*Rosa26*^tdTomato/+^ kidneys (*n* = 2), immunolabelled for PBX1 and tdTomato after E9.5 tamoxifen, showing tdTomato⁺ PBX1⁺ nuclei (arrowheads) in renal stroma. Scale bar: 50 μm. (**d**) Left: Yolk sac from E11.5 *Osr1*^eGFP-CreERT2/+^;*Rosa26*^tdTomato/+^ embryos labelled for EMCN, LYVE1 and tdTomato following E8.5 induction. Scale: 200 μm. Right: Higher magnification of boxed region shows LYVE1⁺ EMCN⁺ vessels (asterisks) lacking tdTomato labelling. Scale bar: 30 μm.

**Figure S6.**
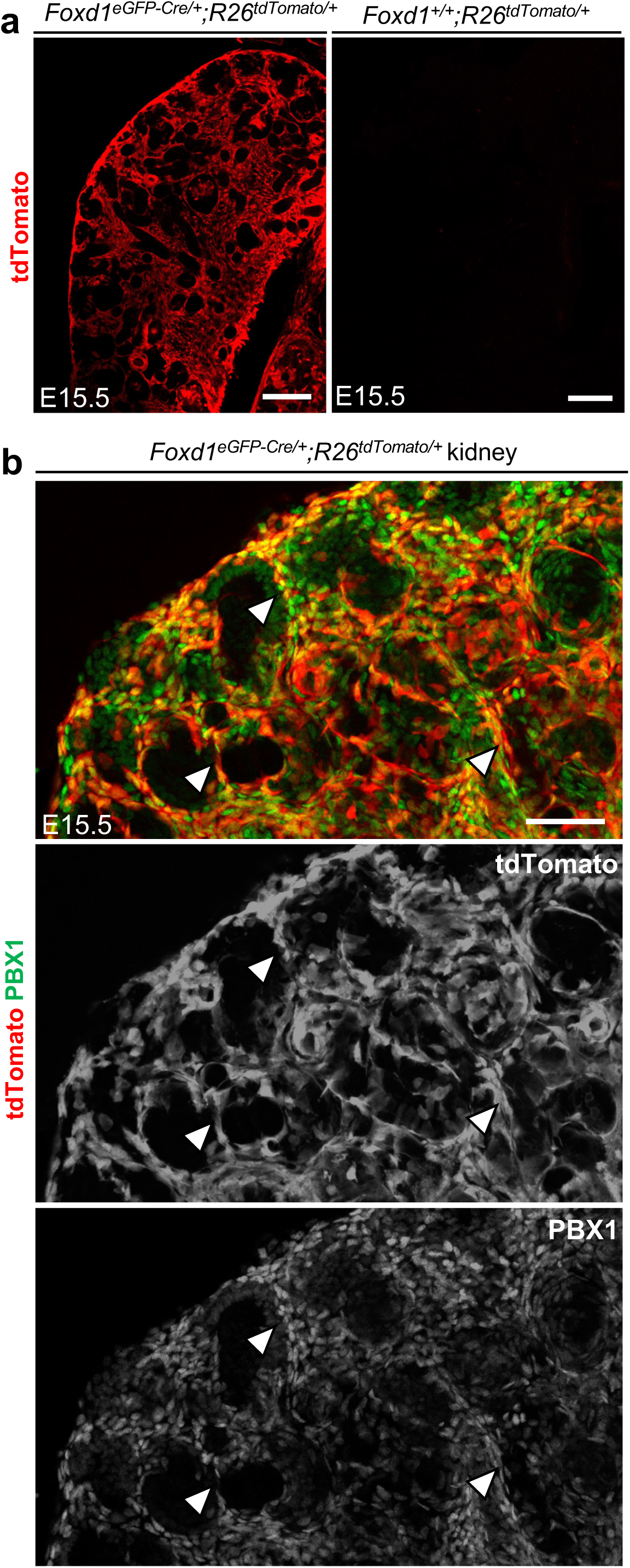
Validation of *Foxd1*^+^ progenitor lineage labelling in the kidney. (**a**) Representative images of E15.5 *Foxd1*^eGFP-Cre/+^;*Rosa26*^tdTomato/+^ kidneys (*n* = 2), immunolabelled for tdTomato (left) and compared with Cre-negative littermates (right). tdTomato expression is seen in a stromal pattern. Scale bar: 100 μm. (**b**) High-resolution confocal z-projection of *Foxd1*^eGFP-Cre/+^;*Rosa26*^tdTomato/+^ kidneys immunolabelled for PBX1 and tdTomato. In the cortex, tdTomato⁺ signal colocalises with PBX1⁺ stromal nuclei (arrowheads). Scale bar: 50 μm.

**Figure S7.**
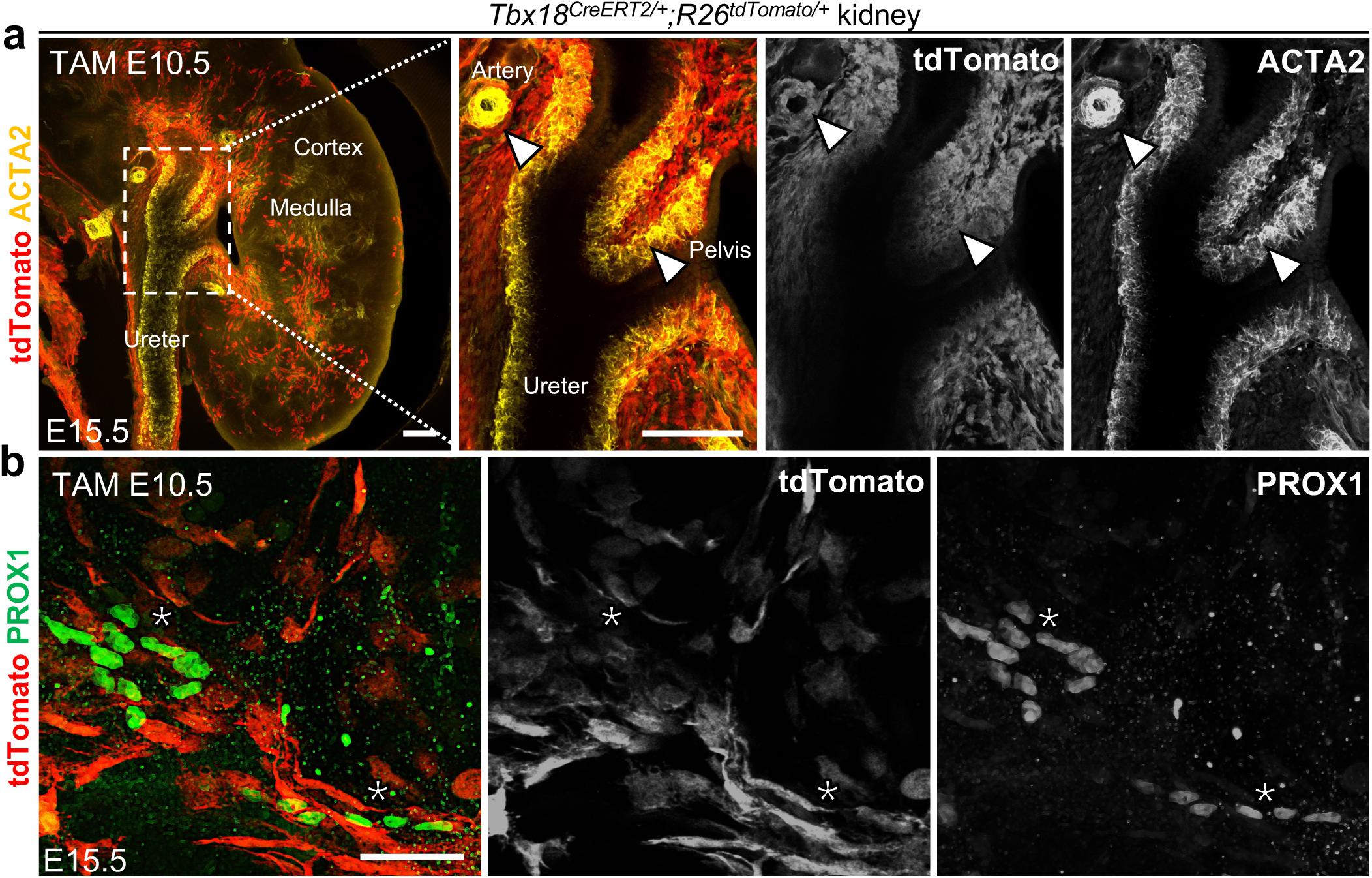
*Tbx18*^+^ progenitor lineage labelling in the kidney shows no contribution to lymphatics. (**a**) Left: Representative low-resolution z-projection of E15.5 *Tbx18*^CreERT2/+^;*Rosa26*^tdTomato/+^ kidneys (*n* = 3), immunolabelled for tdTomato and the smooth muscle, ACTA2. Tamoxifen induction was performed at E10.5. The ureter, medulla and cortex are demarcated. Scale bar: 100 μm. Right: magnified dashed box region from left panel, showing co-expression of tdTomato and ACTA2 within ureteric smooth muscle and vascular smooth muscle (asterisk). Scale bar: 30 μm. (**b**) High-resolution confocal z-projection of the kidney hilum from *Tbx18*^CreERT2/+^;*Rosa26*^tdTomato/+^ kidneys, immunolabelled for PROX1 and tdTomato. In the kidney hilum, no co-expression of PROX1 and tdTomato is observed. Scale bar: 30 μm.

**Figure S8.**
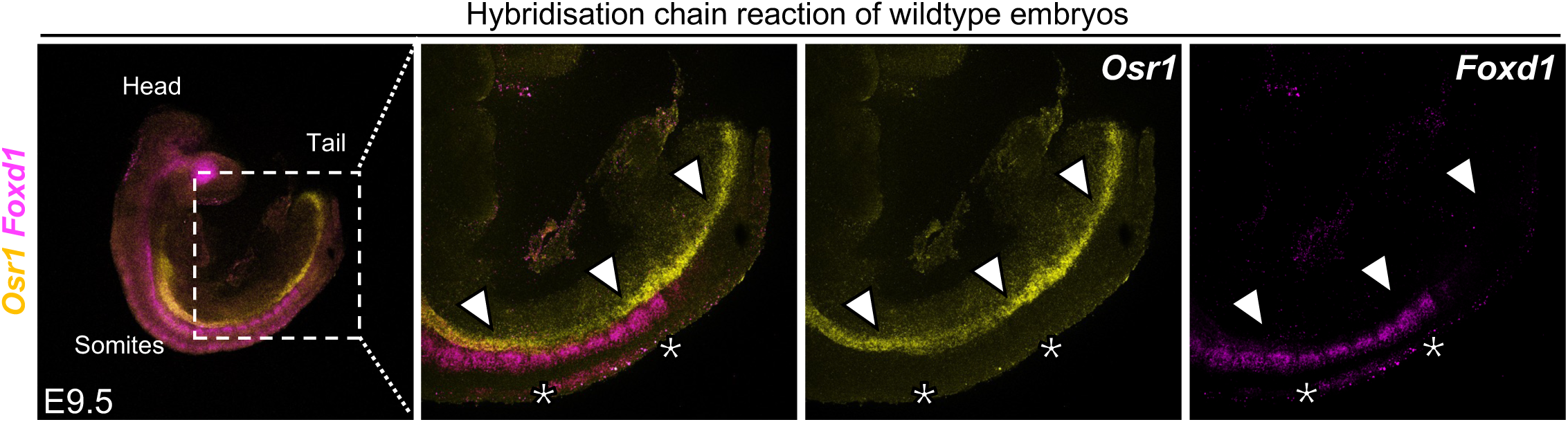
Early expression of *Foxd1*^+^ within somites. Left: Wholemount HCR showing *Osr1* and *Foxd1* expression at E9.5, representative of three embryos pooled across two litters. The head, tail and somites are demarcated. Right: High magnification of boxed region from left panel. *Osr1*⁺ intermediate mesoderm (arrowheads) is distinct from *Foxd1*⁺ cells (asterisks), the latter being expressed within somites. Scale bars: 200 μm.

**Figure S9.**
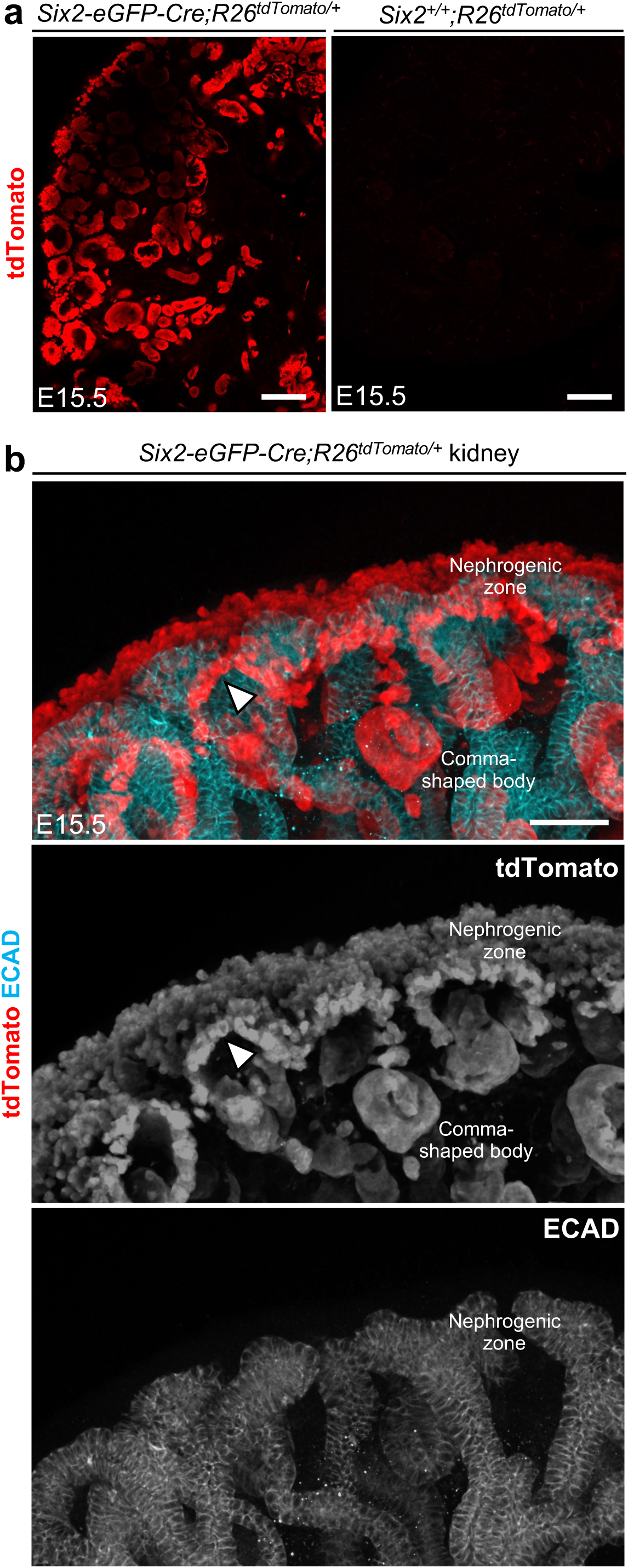
Validation of *Six2*^+^ progenitor lineage labelling in the kidney. (**a**) Representative images of E15.5 *Six2-eGFP-Cre;Rosa26*^tdTomato/+^ kidneys (*n* = 2), immunolabelled for tdTomato (left), compared with littermate controls lacking Cre (right). tdTomato expression is observed in an epithelial pattern. Scale bar: 100 μm. (**b**) High-resolution confocal z-projection of *Six2-eGFP-Cre;Rosa26*^tdTomato/+^ kidneys immunolabelled for ECAD and tdTomato. Strong tdTomato signal is detected in the nephrogenic zone, ECAD⁺ comma-shaped bodies, and cap mesenchyme (arrowhead). Scale bar: 50 μm.

## METHODS

### Mouse husbandry

Mouse experiments were approved across four university sites. Experiments at UK sites (UCL Great Ormond Street Institute of Child Health and UCL Institute of Ophthalmology) were UK Home Office approved and performed in accordance with the Animals (Scientific Procedures) Act 1986. Experiments at SickKids Hospital Research Facility were approved by the Animal Ethics Committee and were carried out in accordance with the regulations of the Canadian Council on Animal Care. Experiments at the University of Colorado were approved by AALAC and procedures were performed in accordance with animal protocols approved by the IACUC at the Anschutz Medical Campus. Mice were kept in specific pathogen-free conditions with unrestricted access to standard chow and water. Ear biopsies were collected at 2–3 weeks of age and stored at –20°C for genotyping. All mice were maintained on a C57Bl/6 background. The following strains were used: *Osr1^eGFP-CreERT2^,* (MGI:3827104, maintained at UCL Great Ormond Street Institute of Child Health), *Six2-eGFP-Cre* (MGI:3848345, maintained at the SickKids Hospital Research Facility), *Foxd1^eGFP-Cre^* (MGI:4359653, maintained at SickKids Hospital Research Facility), *Foxd1^eGFP-CreERT2^* (MGI:J:158051, maintained at UCL Great Ormond Street Institute of Child Health), *Tbx18^CreERT2^*(MGI:J:253913, maintained at the University of Colorado), *Tie2-Cre* (MGI:2450311, maintained at UCL Great Ormond Street Institute of Child Health), *Kit^CreERT2^*(MGI:5543260, maintained at UCL Institute of Ophthalmology)*, Apln^CreERT2^* (MGI:5637737, maintained at UCL Great Ormond Street Institute of Child Health) and *R26^tdTomato^* (MGI:J:155793, maintained at each of the research sites). To obtain embryos, females were time-mated, with the morning of plug detection designated as E0.5. Males carrying Cre or CreERT2 (heterozygous) were crossed with *R26^tdTomato/tdTomato^* females, except for *Apln^CreERT2^* crosses, in which CreERT2 was inherited maternally.

### Activation of CreERT2

Pregnant dams carrying CreERT2-expressing embryos were injected intraperitoneally with either tamoxifen (≥99% purity, T5648, Sigma-Aldrich) or 4-hydroxytamoxifen (≥70% Z-isomer, H6278, Sigma-Aldrich). Compounds were dissolved in ethanol and diluted to 10 mg/ml in peanut oil (P2144, Sigma-Aldrich), with the exception of *Tbx18^CreERT2^* crosses, in which case tamoxifen was dissolved in corn oil (C8267, Sigma-Aldrich) at a concentration of 20mg/ml. Injections were performed at midday using 27G hypodermic needles (Becton Dickinson).

### Genotyping

Cre genotyping was performed using the following primers: Forward: TGGAAAATGCTTCTGTCCGTTTGC; Reverse: AACGAACCTGGTCGAAATCAGTG. PCR was performed using the KAPA2G Fast HotStart genotyping mix (Merck), with a touchdown protocol. Products were resolved via 2% agarose gel electrophoresis.

### Hybridisation chain reaction

Wild-type embryos at various stages were collected from at least three independent litters, fixed overnight in 4% paraformaldehyde (PFA), and processed for whole-mount HCR. Samples were dehydrated in graded methanol (MeOH)/PBS-Tween (0.1%) series. HCR was performed as previously described^83^. DNA probe sets targeting *Osr1, Pax3*, and *Foxd1* transcripts, along with staining reagents and hairpin amplifiers, were supplied by Molecular Instruments. Embryos were stained with DAPI and mounted in CitiFluor AF1 on custom chamber slides.

### Wholemount immunofluorescence and optical clearing

Unless otherwise stated, all reagents were from Sigma Aldrich. Embryonic kidneys and other tissues were dissected in PBS, fixed in 4% PFA overnight at 4°C, washed and stored in PBS with 0.02% sodium azide. Kidneys were sectioned into 250-300 μm-thick slices using a VT1200S Automated Vibrating Microtome (Leica Microsystems). Samples were cleared and stained using a modified iDISCO+ protocol^84^, as previously described^29,48^. Primary antibodies used were: rat anti-EMCN monoclonal (1:50, clone: V.5C7, sc-53941, Santa Cruz), goat anti-PROX1 polyclonal (1:200, AF2727, R&D Systems), rabbit anti-PROX1 polyclonal (1:200, ABN278, Merck), hamster anti-PDPN monoclonal (1:100, clone 8.1.1, 14-5381-82, ThermoFisher Scientific), rat anti-PECAM1 monoclonal (1:50, clone: MEC 13.3, 550274, BD Biosciences) rabbit anti-ACTA2 polyclonal (1:50, ab5694, Abcam), rabbit anti-RFP polyclonal (1:100, 600-401-379, Rockland), goat anti-RFP polyclonal (1:100, AB1140, SicGen), rabbit anti-PBX1 polyclonal (1:100, 4342S, Cell Signalling Technology), goat anti-LYVE1 polyclonal (1:100, AF2089 Abcam), rabbit anti-ECAD monoclonal (1:50, clone: #3195, 24E10, Cell Signalling Technology). Secondary antibodies were all AlexaFluor fluorescent-conjugated, used at 1:200 with excitations at 488 nm, 546 nm, 633 nm or 647 nm. Optical clearing was performed by dehydrating in a methanol series and incubation in benzyl alcohol:benzyl benzoate (BABB) in a 1:2 ratio. Cleared tissues were mounted in BABB on glass FluoroDishes (VWI International) and inverted immediately prior to imaging.

### Confocal microscopy

Images were acquired on an LSM880 upright confocal microscope (Carl Zeiss) using 5×/NA 0.16 EC Plan-Neofluar and 10×/NA 0.5 W-Plan Apochromat water-dipping objectives. Detection was via multi-alkali and GaAsP PMTs or Airyscan Fast (4Y). Laser lines included 488, 561, and 633 nm. Imaging for *Six2-eGFP-Cre* or *Foxd1^eGFP-Cre^* lines was performed at the SickKids Imaging Facility using a Leica SP8 Lightning confocal microscope with a 20×/0.75 objective and 4× HyD detectors with resonant scanning.

### Image processing

For 3D imaging data, donfocal z-stacks were imported into Imaris (v8, Bitplane) for 3D reconstruction and isosurface rendering of tdTomato fluorescence. For HCR datasets, FIJI was used for image processing and maximum intensity projection was performed of representative embryo halves.

### Quantification of lymphatic labelling

Quantification of LECs was performed manually in FIJI. For each kidney, ≥3 randomly selected, non-overlapping volumes were analysed from 300 µm slices. PROX1⁺ nuclei were manually counted and assessed for tdTomato co-labelling using the Multi-point tool. Data were pooled by kidney and mean labelling percentages calculated.

### Analysis of single-nucleus RNA sequencing data

scRNAseq data were previously published^55^. Gene expression plots were generated from the interactive browser https://omg.gs.washington.edu/ using the ‘kidney development’ module, and relevant images exported.

### Reproducibility, statistics and data presentation

All experiments included at least two kidneys from different embryos per litter and were replicated across two or more independent litters. Exact *n* values are provided in the manuscript and corresponding figure legends. Samples were pooled and analysed randomly by litter. To compare lineage-labelling proportions, data distributions were assessed for normality and variance equality; all groups passed these criteria, allowing for parametric testing. Two-group comparisons were made using unpaired two-tailed Student’s *t*-tests; comparisons across multiple groups used one-way ANOVA with Tukey’s post hoc test. Data are presented as mean ± standard error of the mean (SEM), with error bars indicating SEM. A *p* value ≤ 0.05 was considered statistically significant.

## Data availability

All imaging files are available by contacting the lead author upon reasonable request. snRNA-seq data is available via the original publication^55^.

## Acknowledgements

The authors acknowledge Dr Peter Baluk (University of California San Francisco), Professor Juan Pedro Martinez-Barbera (University College London), Dr Robert D’Cruz and Dr Kimberley Lau (SickKids Hospital) for valuable discussions and technical expertise. This work was supported by a Wellcome Trust Investigator Award (220895/Z/20/Z) to DAL, alongside grants from Kidney Research UK (IN_012_20190306), a Rosetrees Trust PhD Plus Award (PhD2020\100012), a Foulkes Foundation Fellowship, and a Wellcome Trust Accelerator Award (314710/Z/24/Z) to DJJ. DJJ is also supported by the Specialised Foundation Programme at the East of England NHS Deanery. ASW acknowledges grant support from the Medical Research Council (MRC)-National Institute for Health and Care Research (NIHR) Rare Disease Research Platform MR/Y008340/1 and MRC Project Grant APP14742. DAL’s laboratory is supported by the NIHR Biomedical Research Centre at Great Ormond Street Hospital for Children NHS Foundation Trust and University College London.

## Author contributions

DJJ, PS, PR, NDR and DAL conceived the study design with support from ASW and CR. DJJ, LGR, AS designed experiments, which were performed and analysed together with CR, MKJ, SI, LAR, CR, JCC and JS. DM assisted with confocal microscopy and 3D imaging protocols. DJJ and DAL wrote the first draft of the manuscript, which was subsequently revised by all others, who approved the final version. DJJ, LGR and AS contributed equally to this work. DAL is the corresponding author.

## Materials & Correspondence

Correspondence and materials request should be addressed to DJJ or DAL.

